# SSU72 phosphatase is a telomere replication terminator

**DOI:** 10.1101/382564

**Authors:** Jose Miguel Escandell, Edison S. Mascarenhas Carvalho, Maria Gallo-Fernandez, Clara C. Reis, Samah Matmati, Inês Matias Luís, Isabel A. Abreu, Stéphane Coulon, Miguel Godinho Ferreira

## Abstract

Telomeres, the protective ends of eukaryotic chromosomes, are replicated through concerted actions by conventional DNA polymerases and telomerase, though the regulation of this process is not fully understood. Telomere replication requires (C)-Stn1-Ten1, a telomere ssDNA-binding complex that is homologous to RPA. Here, we show that the evolutionarily conserved phosphatase Ssu72 is responsible for terminating the cycle of telomere replication in fission yeast. Ssu72 controls the recruitment of Stn1 to telomeres by regulating Stn1 phosphorylation at S74, a residue that lies within the conserved OB fold domain. Consequently, ssu72Δ mutants are defective in telomere replication and exhibit long 3’ overhangs, which are indicative of defective lagging strand DNA synthesis. We also show that hSSU72 regulates telomerase activation in human cells by controlling the recruitment of hSTN1 to telomeres. Thus, in this study, we demonstrate a previously unknown yet conserved role for the phosphatase SSU72, whereby this enzyme controls telomere homeostasis by activating lagging strand DNA synthesis, thus terminating the cycle of telomere replication.

## Introduction

Telomeres are protein-DNA complexes that form the ends of eukaryotic chromosomes (reviewed in ^1^). Telomeres predominantly function to prevent the loss of genetic information and to inhibit DNA repair at the chromosome termini, thus maintaining telomere protection and genome stability. Loss of telomere regulation has been linked to two main hallmarks of cancer: replicative immortality and genome instability ^2^. Telomeres face an additional challenge: DNA replication. Due to G-rich repetitive DNA sequence and protective structures, telomeres represent a natural obstacle for passing replication forks ^3^. Replication fork collapse can lead to the loss of whole telomere tracts. To counteract these effects, telomerase (Trt1 in S. *pombe* and TERT in mammals) is responsible for adding specific repetitive sequences to telomeres, compensating for the cell’s inability to replicate chromosome ends ^4^. However, it is currently not understood how telomerase activity is regulated and how the telomerase cycle is coupled to telomeric DNA replication. Intriguingly, several DNA replication proteins are required for proper telomere elongation ^5^. Conversely, specific telomere components are themselves required for proper telomere replication and telomere length regulation^6,7^, suggesting that there is a very thin line separating telomere replication and telomerase activity.

Using fission yeast, Chang et al. proposed a dynamic model that demonstrates how telomere replication controls telomere length and how this is carried out by the telomere complex (Chang et al., 2013). The telomere double strand components Taz1, Rap1, and Poz1 promote the recruitment of Polα-Primase to telomeres. Because shorter telomeres possess less Taz1/Rap1/Poz1, Polα-Primase recruitment, and therefore lagging-strand synthesis, is delayed at chromosome ends, leading to the accumulation of ssDNA at telomeres. This event results in the activation of the major checkpoint protein kinase Rad3^ATR^ and the subsequent phosphorylation of telomeric Ccq1-T93, a step required for telomerase activation. Thus, as a consequence of delayed Polα-Primase recruitment to short telomeres and the subsequent accumulation of ssDNA, Rad3^ATR^ is transiently activated leading to telomerase recruitment and telomere elongation.

Another complex known as CST (Cdc13/Stn1/Ten1 in S. *cerevisiae* and CTC1/STN1/TEN1 in mammals), is known to control telomere replication. This complex is responsible for both protection from 5’ strand nucleolytic degradation and recruitment of the Polα-primase complex to telomeres, thus promoting telomere lagging-strand DNA synthesis (Grossi et al., 2004; Lin and Zakian, 1996). Notably, CST is not only required to recruit Polα-primase but is also responsible for the switch from primase to polymerase activity, which is required for gap-less DNA replication ^11^. In humans, in addition to its role in telomere replication ^12^, the CST complex also functions as a telomerase activity terminator ^13^ by inhibiting telomerase activity through primer confiscation and direct interaction with the POT1-TPP1 dimer. However, the mechanism regulating these CST functions remains unknown. In fission yeast, although *stn1+* and *ten1+* homologs exist, no Cdc13/CTC1 homolog has been found to date ^14^. Recent studies have revealed that Stn1 is required for telomere and subtelomere replication ^15^ and ^16^, supporting the conserved role of fission yeast (C)ST in DNA replication.

In agreement with the replication model proposed by ^17^ and reviewed in ^18^, the telomere-binding protein, Rif1 was shown to regulate telomere DNA replication timing by recruiting Glc7 phosphatase to origins of replication and inhibiting Cdc7 activities in budding yeast (Hiraga et al., 2014; Mattarocci et al., 2014). Notably, this role is conserved in other organisms such as fission yeast ^22^ and human cells ^23^. Importantly, *rif1* mutants display long telomeres; this effect is suggested to be a result of origin firing dysregulation ^18^. However, how telomere replication is terminated and how this is coupled with the regulation of telomere length remains unknown. Here, we report that the phosphatase family member Ssu72 displays a conserved role as a telomere replication terminator. Ssu72 was previously identified as an RNA polymerase II C-terminal domain phosphatase and is highly conserved from yeast to human ^24^. In addition, Ssu72 functions as a cohesin-binding factor involved in sister-chromatid cohesion by counteracting the phosphorylation of SA2, another cohesion complex member ^25^. In fission yeast, in addition to regulating RNA polymerase activity, Ssu72 has recently been shown to regulate chromosome condensation ^26^. However, none of the previous studies have noted deregulated telomere replication. Our data strongly support an unexpected role for Ssu72 in controlling lagging-strand synthesis through the regulation of Stn1 Serine74 phosphorylation, thus reducing telomeric ssDNA and inhibiting telomerase recruitment.

## Results

### Ssu72 is a negative regulator of telomere elongation

We carried out a genome-wide screen for regulators of telomere homeostasis in *S. pombe* using a commercially available whole-genome deletion library (*Bioneer* corporation). This library allowed us to identify new non-essential genes involved in telomere homeostasis in fission yeast (**Figure 1A**). Of the genes identified from the screen, we selected the highly conserved phosphatase *ssu72+* (SPAC3G9.04) as the most promising candidate for further characterization. We generated a deletion mutant (*ssu72*Δ) as well as a point mutant devoid of phosphatase activity (*ssu72-C13S*) and found that these two mutants possess longer telomeres (**Figure 1B**). Additionally, we found that Ssu72 localized to telomeres in a cell cycle-dependent manner. We performed cell cycle synchronization using a *cdc25-22* block-release method in a *ssu72-myc* tagged strain and measured Ssu72 binding to telomeres by chromatin immunoprecipitation (ChIP). Cell cycle phases and synchronization efficiency were measured using the cell septation index. Ssu72-myc is recruited to telomeres in late S phase and declines later in the cell cycle (**Figure 1C**). Interestingly, Ssu72 is recruited to telomeres at approximately the same time as the arrival of the lagging strand machinery at chromosome ends ^17^.

**Figure 1.**
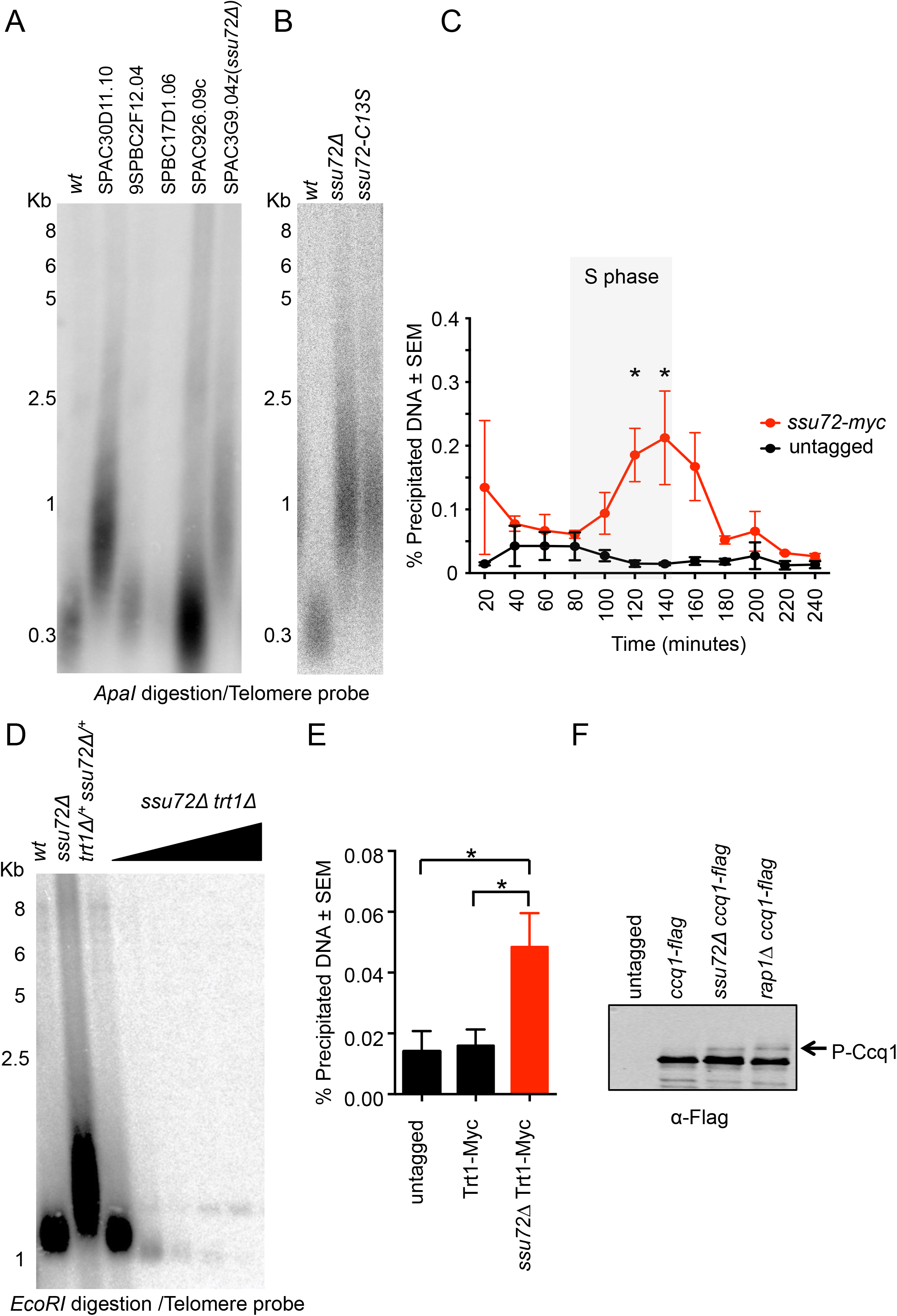
Genetic screen identifies Ssu72 as telomerase regulator. A) We identified previously unknown telomere regulators in fission yeast using the haploid *S. pombe* whole-genome gene deletion library including *ssu72+* (SPAC3G9.04). B) Telomere length in *wt, ssu72*Δ and *ssu72-C13S* (point mutant on the phosphatase active site) strains were measured by Southern Blot in *ApaI* digested samples using a telomeric probe. C) Ssu72 recruitment to telomeres is cell cycle regulated. Ssu72 was myc-tagged in a *cdc25^ts^* strain and ChIP analysis was carried out in cell cycle synchronized populations. Septa formation was used as readout for S-phase. n > 3; *p <0.05 based on a two-tailed Student’s *t*-test to *ssu72+* control samples. Error bars represent standard error of the mean (SEM). D) Telomere length of *ssu72*Δ is dependent on telomerase. Diploid strains with the appropriate phenotypes were sporulated and double mutants *trt1Δ ssu72*Δ were streaked for multiple passages (triangle indicates increased number of generations). E) Telomerase is recruited to telomeres in the absence of Ssu72. ChIP analysis for Trt1-myc in *wt* and *ssu72*Δ was performed as described in material and methods using a non-tagged strain as a control. n > 3; *p <0.05 based on a two-tailed Student’s *t*-test to control sample. Error bars represent standard error of the mean (SEM). F) The telomerase activator Ccq1 is phosphorylated in *ssu72*Δ cells. *rap1*Δ cells were used as positive control. Western blots were performed using Ccq1-flag tagged strains.

*ssu72*Δ cells displayed increased (~1 Kb) telomere lengths compared to wild-type telomeres (~300 bp) (**Figure 1B**). We set out to understand the nature of telomere elongation in the *ssu72* mutant background. To test if the telomere elongation was dependent on telomerase, *trt1*Δ (deletion mutant for the catalytic subunit of telomerase) and *ssu72*Δ double heterozygous diploids were sporulated. Of the resulting tetrads, *trtl*Δ and *trtlΔ ssu72*Δ double mutants were selected and streaked for several generations in order to facilitate telomere shortening in the absence of telomerase. While *ssu72* mutant cells displayed long telomeres, *ssu72Δ trt1*Δ double mutant and trt1Δ single mutant cells displayed similarly shortened telomeres (**Figure 1D**). ChIP experiments consistently demonstrated an accumulation of Trt1-myc at *ssu72*Δ telomeres compared to wt cells (**Figure 1E**). Thus, the longer telomeres exhibited by *ssu72*Δ mutants were a consequence of telomerase deregulation.

Two independent studies ^27,28^ showed that Ccq1 phosphorylation at Thr93 is required for telomerase-mediated telomere elongation in fission yeast. Using Western blot shift analysis, we observed that Ccq1 was phosphorylated in *ssu72*Δ cells when we compared to those of WT strains (**Figure 1F**). To further confirm that telomere elongation was telomerase-dependent, we repeated the previous experiment using a phosphorylation-resistant mutant version of Ccq1 ^27^. We germinated a double heterozygous *ccq1-T93A/+ ssu72Δ/+* mutant and analyzed its progeny. As expected, *ccq1-T93A ssu72*Δ double mutants displayed a similar telomere-shortening rate to that of the *ccq1-T93A* single mutants (**Figure S1**). In agreement with these results, we further showed that telomere length in *ssu72*Δ mutants was dependent on Rad3, the kinase responsible for Ccq1-T93 phosphorylation (**Figure S2A**), and not dependent on the checkpoint kinase Chk1 (**Figure S2B**). In addition, *ssu72Δ rad51*Δ double mutants displayed similar telomere lengths to *ssu72*Δ single mutants (**Figure S2C**). Taken together, our results demonstrate that Ssu72 is a negative regulator of telomerase, possibly counteracting Rad3 activation and Ccq1 phosphorylation.

### Ssu72 phosphatase function is independent of Rif1 and Taz1/Rap1/Poz1

In fission yeast, the presence of telomeric ssDNA results in Rad3 activation and telomere elongation ^27^. Thus, we investigated whether *ssu72*Δ mutants accumulated telomeric ssDNA. We carried out in-gel hybridization assays using a C-rich probe to measure the accumulation of G-rich DNA at telomeres. Notably, the *ssu72*Δ mutant strain showed an almost 6-fold increase in G-rich telomere sequences (**Figure 2A**). We observed that the accumulation of ssDNA at telomeres is increased in *ssu72*Δ mutants compared to *rif1*Δ mutants, though both strains have similar telomere lengths. Further, we consistently detected Rad11^RPA^-GFP localization at telomeres, as measured by live imaging in *ssu72*Δ mutant cells (**Figure 2B**).

**Figure 2.**
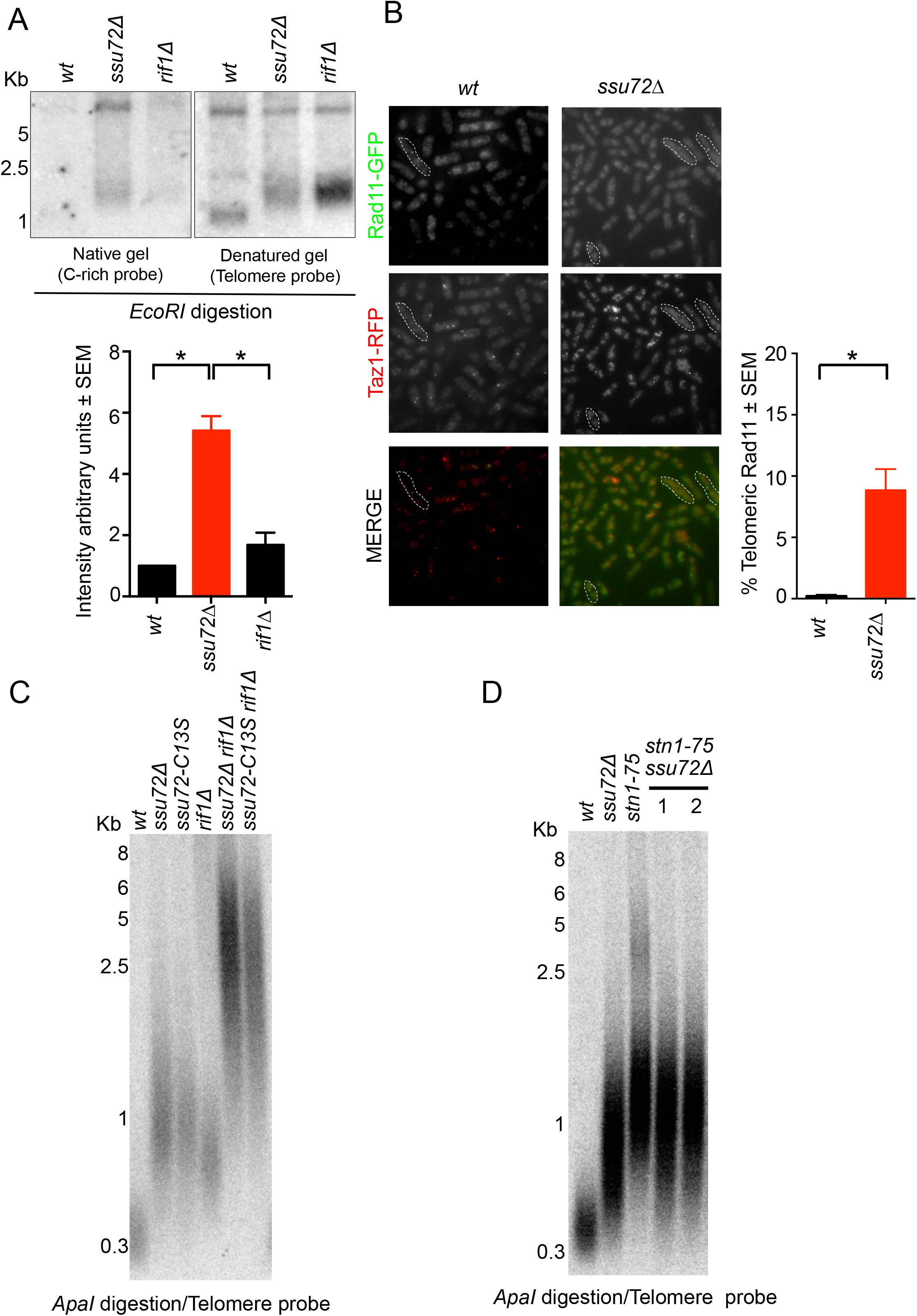
Ssu72 is required for telomeric C-strand. A) *ssu72*Δ telomeres present longer G-rich overhangs than *wt* and *rif1*Δ telomeres. In-gel hybridization in native and denaturing conditions was labelled with a radiolabelled C-rich telomere probe and quantified for ssDNA at the telomeres. n = 2; *p ≤0.05 based on a two-tailed Student’s *t*-test to control sample. Error bars represent Standard error of the mean (SEM). B) RPA (Rad11-GFP) is enriched at *ssu72*Δ telomeres. Colocalization of Rad11-GFP with Taz1-mCherry, used as a telomere marker, was performed in *wt* and *ssu72*Δ cells; n =3; *p ≤0.05 based on a two-tailed Student’s *t*-test to control sample. Error bars represent standard error of the mean (SEM). More than 1000 cells were analyzed in each phenotype C) *ssu72^+^* controls telomere length independently of *rif1^+^*. Epistasis analysis of telomere length of *ssu72*Δ and *ssu72-C13S* (catalytically inactive mutant) with *rif1*Δ was performed by Southern blotting of *Apal* digested DNA using a telomeric probe. D) *ssu72^+^* and *stn1+* regulate telomere length in the same genetic pathway. Epistasis analysis of *ssu72*Δ and *stn1-75* performed by Southern blotting of *Apal* digested DNA using a telomeric probe. Two independently generated *ssu72Δ stn1-75* double mutants are shown.

Recently, the telomere-binding protein Rif1 was found to control DNA resection and origin firing by recruiting PP1A phosphatase to double strand breaks and origins of replication ^20-22,29^. We wondered if Rif1 was also responsible for recruiting the Ssu72 phosphatase to telomeres. To test this hypothesis, we combined *ssu72*Δ and *ssu72-C13S* (catalytically dead) mutants with *rif1*Δ and carried out of telomere length epistasis analyses. While single mutants displayed telomere lengths of 1 Kb, *ssu72Δ rif1*Δ and *ssu72-C13S rif1*Δ double mutants displayed telomeres that were longer than 3 Kb (**Figure 2C**). Thus, our data show that Rif1-mediated regulation of telomere length is independent of Ssu72 in fission yeast.

### Ssu72 controls telomere length through the Stn1-Ten1 complex

A second highly conserved protein complex regulates telomere length and telomerase activity. The budding yeast CST complex (Cdc13^CTC1^, Stn1 and Ten1) plays opposing roles at the telomeres. Cdc13 is required for telomerase recruitment and is activated through its interaction with Est1, a subunit of telomerase ^30^. This interaction is promoted by the phosphorylation of Cdc13 at T308 by Cdk1(Cdc28). In contrast, the Siz1/2-mediated SUMOylation of Cdc13 at Lys908 promotes its interaction with Stn1 ^31^. This interaction is required, with Ten1, for polymerase alpha complex recruitment and telomere lagging-strand DNA synthesis ^8^. However, the regulatory mechanism underlying these two opposite functions remains unknown. Despite the lack of Cdc13^CTC1^ homologs in fission yeast, the Stn1-Ten1 complex appears to play similar roles to those found in budding yeast and mammals (reviewed in ^32^). Consequently, we hypothesized that Ssu72 controls telomere length through the Stn1-Ten1 complex. Because fission yeast *stn1* and *ten1* deletion mutants lose telomeres completely and survive only with circular chromosomes ^14^, we carried out our experiments in mutants carrying a hypomorphic *stn1-75* allele ^33^. Similar to *ssu72*Δ mutants, *stn1-75* mutants possess long telomeres (~1 Kb) (**Figure 2D**). In contrast to our previous genetic studies, *stn1-75 ssu72*Δ double mutants displayed similar telomere lengths to those of single mutants. This result suggests that Ssu72 controls telomere length through the same pathway as the Stn1-Ten1 complex.

We then decided to investigate this genetic interaction using a different strategy. Fission yeast Stn1 is recruited to telomeres in a cell cycle-dependent manner ^34,35^, with peak telomere association in the S/G2 phases of the cell cycle. This coincides with Ssu72 recruitment to telomeres, as observed in our synchronization experiments (**Figure 1C**). Given the genetic association, we asked whether the recruitment of Stn1 to telomeres was dependent on the function of Ssu72. To test this hypothesis, we performed Stn1-myc ChIP experiments in *ssu72*Δ cells throughout the cell cycle (**Figure 3A**). As has been previously demonstrated, Stn1-myc was recruited to telomeres in S/G2 cells ^36^. We observed that the recruitment of Stn1 to telomeres was severely impaired in the absence of Ssu72 (**Figure 3A**). Thus, our results indicate that Ssu72 functionality is required for Stn1 recruitment to telomeres.

**Figure 3.**
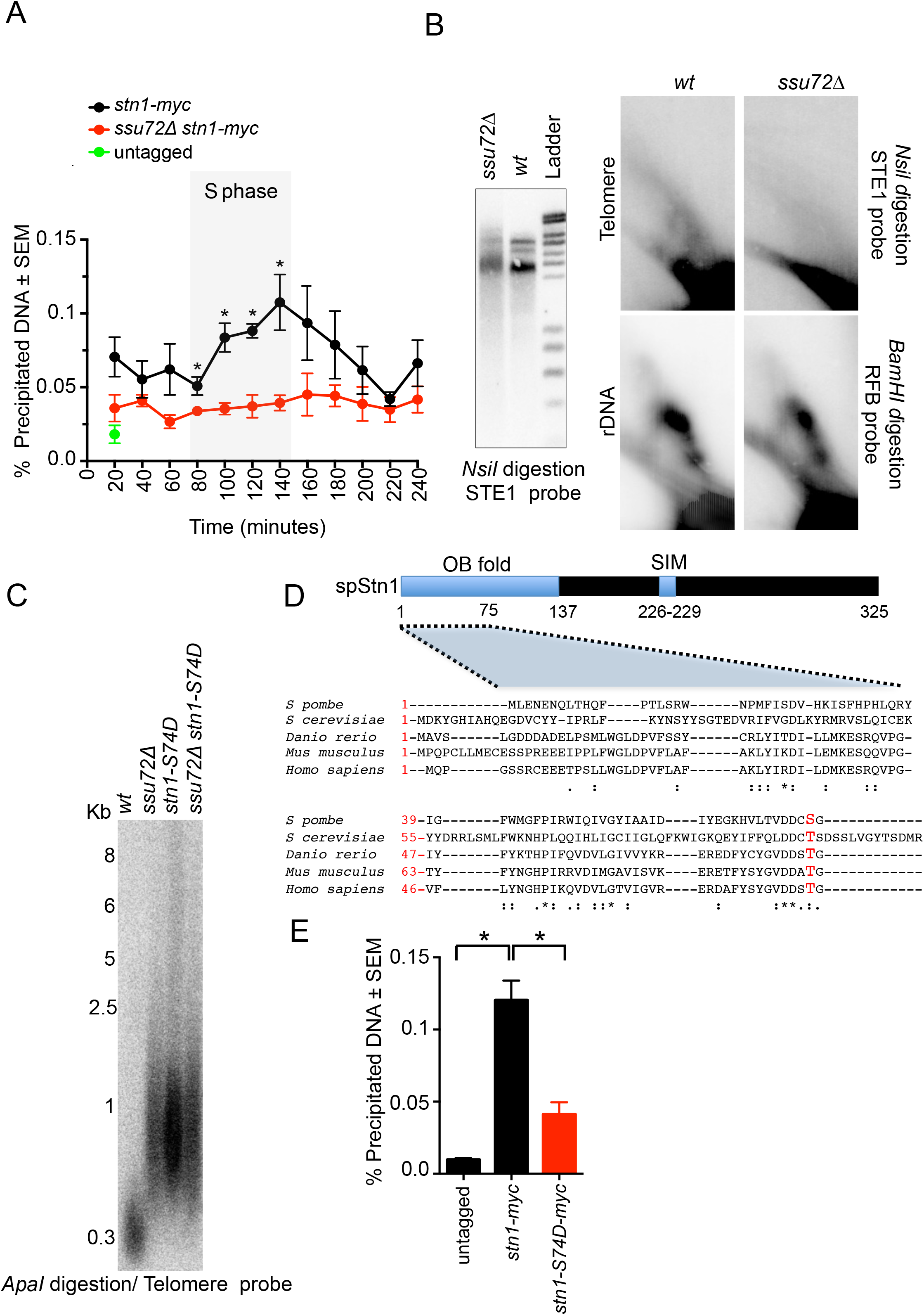
Ssu72 controls Stn1 telomere recruitment and phosphorylation. A) Ssu72 is required for telomere recruitment of Stn1 in late S phase. ChIP analysis of *stn1-myc* in *wt* and *ssu72*Δ cells was performed in synchronized *cdc25^ts^* cells. n > 3; *p <0.05 based on a two-tailed Student’s *t*-test to *ssu72+* control samples. Error bars represent standard error of the mean (SEM). B) 2D-gel analysis of NsiI telomeric fragments of *wt* and *ssu72*Δ strains. Smart ladder from *Eurogentec* was used for DNA size measurement. C) Serine 74 substitution to a phosphomimetic aspartate amino acid (*stn1-S74D*) is sufficient to confer *ssu72*Δ telomere defects. Telomere length epistasis analysis of *ssu72*Δ and *stn1-S74D* mutants were performed by Southern blotting of *Apal* digested genomic DNA using a telomeric probe. D) Sequence alignment of Stn1 highlighting serine 74 identified in fission yeast as a phosphorylated residue. E) Similar to *ssu72*Δ mutants, *stn1-S74D* is defective in telomere recruitment. ChIP analysis of *stn1-myc* and *stn1-S74D-myc* using a non tagged strain as a control. n = 3; *p <0.05 based on a two-tailed Student’s *t*-test to control sample. Error bars represent standard error of the mean (SEM).

Based on these findings, we asked whether DNA replication dynamics were affected at *ssu72*Δ mutant chromosome ends. Genomic DNA derived from *WT* and *ssu72*Δ cells was isolated, subjected to NsiI digestion, and analyzed on 2D-gels. Chromosome ends were revealed by Southern blotting using a telomere-proximal STE1 probe ^6,37^. In the first dimension, we observed three distinct bands for the wt parental strain but only one thick, smeared band for the *ssu72*Δ strain. As expected, we observed Y structures derived from passing replication forks within the *NsiI* fragment in WT cells (Figure 3B). In contrast, Y structures were not observed in *ssu72*Δ mutants. To control for our ability to detect DNA replication in a *ssu72*Δ background, we analyzed the ribosomal DNA replication fork barrier using BamHI-digested genomic DNA probed with an rDNA-specific probe (rDNA-RFB). As expected, replication fork blocks were similarly detected in both WT and *ssu72*Δ mutant strains. Thus, these results indicate that Ssu72 is required for DNA replication at chromosome ends, consistent with the role of Ssu72 in regulating Stn1 recruitment to telomeres.

### Ssu72 regulates Stn1 phosphorylation

Given that Ssu72 phosphatase activity is required to regulate telomere length and that Ssu72 is recruited to telomeres during the S/G2 phases, we hypothesized that Ssu72 might regulate Stn1 phosphorylation in a cell cycle-dependent manner. Previous studies have revealed different Cdk1-dependent phosphorylation sites in Stn1 in budding yeast ^38,39^. However, to date, Stn1 phosphorylation sites have not been identified in species outside of *S. cerevisiae*. Moreover, the phosphorylation sites described for budding yeast are not conserved in *S. pombe*. Thus, we decided to take an unbiased approach using mass spectrometry-based analysis of purified fission yeast Stn1. First, we immunoprecipitated Stn1-myc from cells carrying the *ssu72*Δ deletion. Subsequent analysis of the immunoprecipitated material revealed a phosphorylated peptide corresponding to Stn1 Serine-74 (**Figure S3A**). Notably, this serine is not only conserved in *Schizosaccharomyces* (the fission yeast genus) (**Figure S3B**) and budding yeast but also throughout higher eukaryotes, including humans and mice (**Figure 3D**). Therefore, we decided to mutate the Serine-74 residue to aspartic acid (Stn1-S74D), a phosphomimetic amino acid replacement.

Cells harboring the *stn1-S74D* mutation exhibited long telomeres (~1 Kb) similar to those found in *ssu72*Δ cells (**Figure 3C**). We hypothesized that telomere elongation in the *stn1-S74D* strain was telomerase-dependent. Consistent with this hypothesis, *stn1-S74D trt1*Δ double mutant telomeres become shorter after sequential streaks (**Figure S4A**). Importantly, *stn1-S74D ssu72*Δ double mutants displayed similar telomere lengths to those in *stn1-S74D* single mutants. In addition, we performed ChIP experiments in strains expressing Stn1-S74D-myc in order to analyze the recruitment of Stn1 to telomeres. Similar our observations in mutants lacking *ssu72* phosphatase, Stn1-S74D-myc was not efficiently recruited to telomeres (**Figure 3E**). Taken together, our data suggest that fission yeast Stn1 is phosphorylated at Serine-74 to enable its efficient recruitment to telomeres and, consequently, efficient DNA replication and telomerase regulation.

Given that both Stn1 and Ssu72 are recruited to telomeres in the S/G2 phases of the cell cycle and that Stn1-S74 phosphorylation is required for efficient Stn1 recruitment to telomeres, we wondered if Ssu72 phosphatase could counteract the action of a cell cycle-dependent kinase. Due to the central nature of Cdc2^Cdk1^ in regulating the cell cycle, we mutated *ssu72+* in cells carrying the *cdc2-^M68^* temperature-sensitive mutant allele. At permissive temperatures (25°C), *ssu72Δ cdc2-^M68^* strains exhibited a similar telomere length to *ssu72*Δ single mutants (**Figure S4B**). To inactivate Cdc2 activity, we grew *ssu72Δ cdc2-^M68^* strains at semi-permissive higher temperatures. Our results show that progressive inactivation of Cdc2^Cdk1^in *ssu72Δ cdc2-^M68^* double mutants resulted in a gradual decrease in telomere lengths compared to those in *ssu72*Δ strains. Even though Stn1 Serine-74 does not lie within a conserved CDK consensus site, our data suggest that Cdc2^Cdk1^ activity may counteract Ssu72 phosphatase.

### Ssu72 is required for DNA replication at chromosome ends

Stn1, part of the CST complex, performs many different functions. In humans, it has been proposed to be a terminator of telomerase activity due to its higher affinity for telomeric single stranded DNA formed after telomerase activation ^13^. Further, it has been suggested to promote the restart of stalled replication forks ^40,41^. In fission yeast, the ST complex also exhibits this dual function. First, the binding of this complex at telomeres inhibits telomerase action through an interaction with K242-SUMOylated Tpz1 and the SIM domain of Stn1 ^16,33,35^. Secondly, Stn1 participates in telomere and subtelomere semi-conservative DNA replication ^15,16^.

Budding yeast CST promotes lagging strand synthesis by interacting with the catalytic and B-subunits of the DNA polymerase α-primase complex ^8,30,42^. We thus hypothesized that Ssu72 controls the DNA polymerase α-primase complex at fission yeast telomeres. To test this hypothesis, we carried out epistasis analyses with the catalytic subunit of the polymerase α complex *(pol1+)*. A hypomorphic mutation in this subunit results in longer telomeres in fission yeast due to the formation of 3’ overhangs that sustain telomerase activation ^5^. As expected, while *pol1-13* had telomeres of approximately 1 Kb in length, *pol1-13 ssu72*Δ double mutants had similar telomere lengths to single mutants (**Figure 4A**). In addition, we carried out similar epistasis studies with both RNA primase subunits (Spp1 and Spp2) and observed that *spp1-9 ssu72*Δ and *spp2-9 ssu72*Δ double mutants exhibited similar telomere lengths to those of single mutants (**Figure S5A**). Taken together, these results indicate that the functionality of Ssu72 at telomeres relies on the activity of both DNA polymerase α and RNA primase complexes. Thus, similarly to Stn1, Ssu72 controls lagging strand synthesis at telomeres (**Figure S5A**).

**Figure 4.**
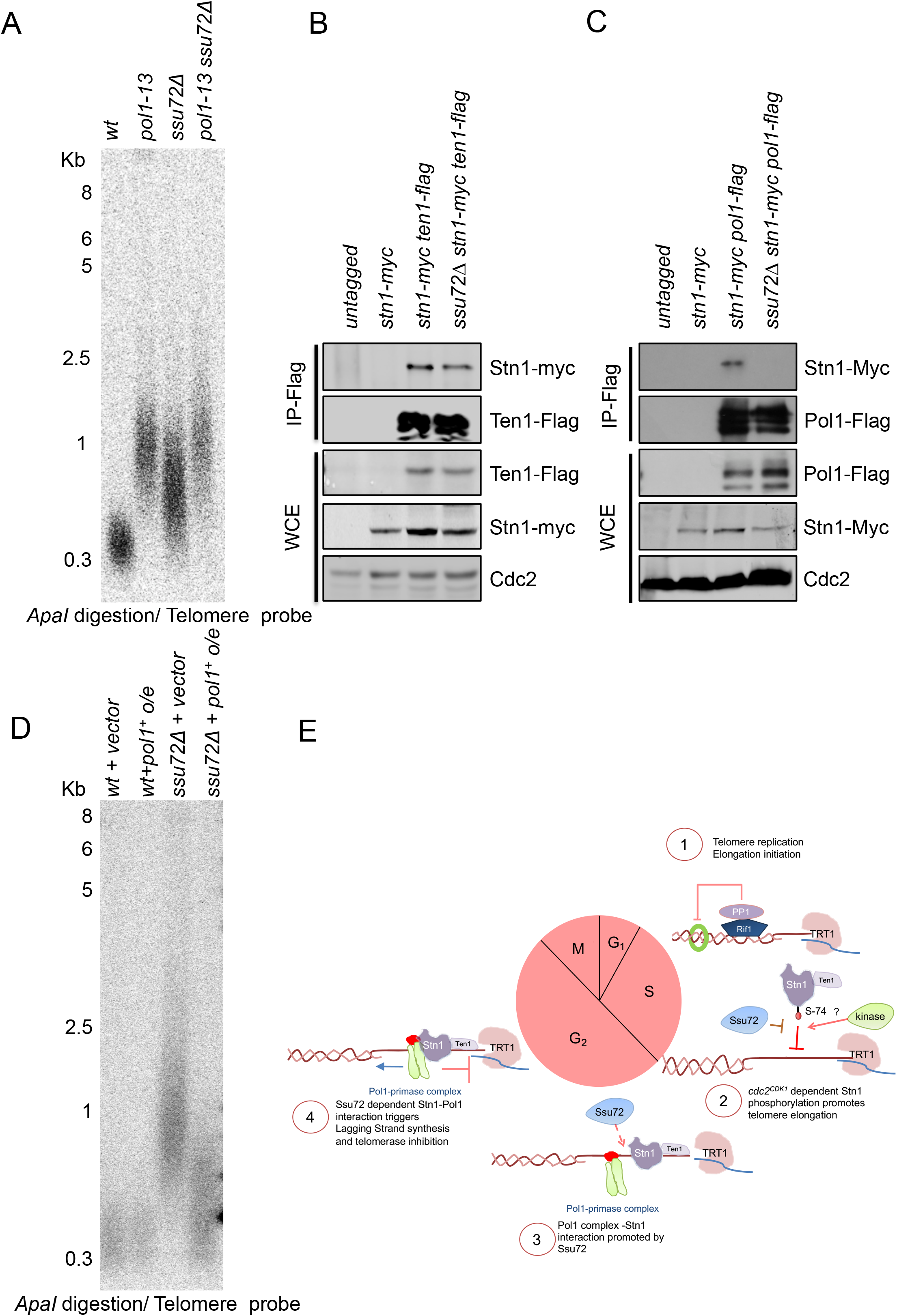
Ssu72 is required for polymerase α activation. A) *ssu72^+^* and DNA polymerase α regulate telomere length in the same genetic pathway. Epistasis analysis of *ssu72*Δ and *pol1-13* performed by Southern blotting of *Apal* digested DNA using a telomeric probe. B) and C) Stn1-Pol1 interaction requires Ssu72. Immunoprecipitation experiments of Pol1-Flag with Stn1-Myc was preformed both in *wt* and *ssu72*Δ mutants. As control we carried out immunoprecipitation experiments of Ten1-Flag with Stn1-Myc in either *wt* or *ssu72*Δ D) overexpression of polymerase alpha rescues telomere defect in *ssu72*Δ cells. Multi-copy vector with polymerase α under thiamine promoter were expressed both in *wt* or *ssu72*Δ cells. E) Proposed model for Ssu72 regulation of telomere replication in fission yeast. See text for details.

We next investigated if Stn1 overexpression was sufficient to rescue the telomere defects observed in *ssu72*Δ mutants. To test this hypothesis, we replaced the *stn1+* endogenous promoter with inducible Thiamine-regulated *nmt* promoters ^43^. We observed that none of the promoters used to overexpress Stn1 rescued the telomere defects of *ssu72*Δ (**Figure S5B**), indicating that Stn1 overexpression was unable to compensate for the defects in *ssu72*Δ.

CST in budding yeast regulates lagging strand synthesis by stimulating DNA polymerase activity through the interaction of Stn1 with Pol1 ^11^. Similarly, we postulated that the mechanism whereby Ssu72 phosphatase controls lagging strand synthesis is achieved by regulating the Stn1-Pol1 interaction. To test this hypothesis, we carried out immunoprecipitation experiments using extracts derived from Pol1-Flag and Stn1-Myc tagged strains. As a control, we first verified that we could purify Ten1-Flag with Stn1-Myc in *ssu72*Δ mutants. We could readily coimmunoprecipitate Ten1-Flag and Stn1-Myc in both the *wt* and *ssu72*Δ strains (**Figure 4B**). Thus, Ssu72 phosphatase does not regulate the Stn1-Ten1 interaction. Similar to what has been observed in budding yeast, we were able to demonstrate the Pol1-Stn1 interaction in these experiments (**Figure 4C**). In contrast, we were unable to immunoprecipitate Stn1-Myc with Pol1-Flag using an anti-Flag antibody in *ssu72*Δ cells. Our results show that Ssu72 is required for the interaction of Stn1 with the polymerase alpha complex, suggesting that Ssu72 functionality relies on Stn1-dependent activation of lagging strand synthesis.

The results of the previous experiment suggested the hypothesis that Ssu72 is required to activate DNA polymerase α at telomeres. To test this, we overexpressed the catalytic subunit of polymerase α *(pol1+)* in cells lacking Ssu72. Previous studies have shown that overexpression of *pol1+* was sufficient to rescue strains with lagging strand synthesis defects ^5^. Remarkably, using *pol1+* expression from multicopy plasmids, we showed that overexpression of *pol1+* in *ssu72*Δ mutants is sufficient to rescue telomere defects (**Figure 4D**). Thus, we propose that Ssu72 phosphatase regulates Stn1 phosphorylation status in order to control Stn1 recruitment to telomeres and DNA polymerase α activation of lagging strand synthesis. We propose a dynamic model where Rif1 dependent phosphatase activities regulate telomere replication initiation by controlling origin firing. Further, we propose that Ssu72 phosphatase functions as a telomere replication terminator by regulating Stn1 recruitment to telomeres in a cell cycle-dependent manner to activate lagging strand synthesis, thus ending the telomere replication cycle (**Figure 4E**).

### SSU72 telomere function is conserved throughout evolution

Because Ssu72 is a highly conserved phosphatase and CST has similar functions in different species, we tested if telomere regulation by SSU72 was conserved in human cells. We were not able to produce human cell lines lacking SSU72 using conventional CRISPR/Cas9 technology, suggesting that SSU72 is essential in humans. In contrast to fission yeast, *SSU72* is an essential gene both in budding yeast ^24,44^ and mice ^45^. Therefore, we decided to use short hairpin RNAs to downregulate SSU72 protein levels. This approach has been previously used in human cells to study the role of SSU72 in mammals ^25^.

Our results show that, similar to fission yeast, knockdown of SSU72 in human cells causes telomere dysfunction. We downregulated SSU72 levels using two specific shRNAs and collected HT1080 cells for analysis of telomere length 6 weeks after infection. In HT1080 cells, the median telomere length is 3.4 Kb in cells transfected with a control Luciferase shRNA (**Figure 5A**). As observed in fission yeast, downregulation of SSU72 using shRNAs against CDS sequence (knockdown efficiency 85 %) or UTR sequence (knockdown efficiency 92 %) results in an increase in telomere length to 3.7 Kb and 3.8 Kb, respectively (**Figure 5A**). The observed telomere elongation results from telomerase activity. To test this, we used the telomerase inhibitor BIBR1532. Treatment of cells with BIBR1532 resulted not only in the inhibition of telomere elongation in shSSU72-infected cells but also in a general decrease in telomere length in all treated cells (**Figure 5A**). Thus, as in fission yeast, SSU72 controls telomere length by regulating telomerase function in human cells.

**Figure 5.**
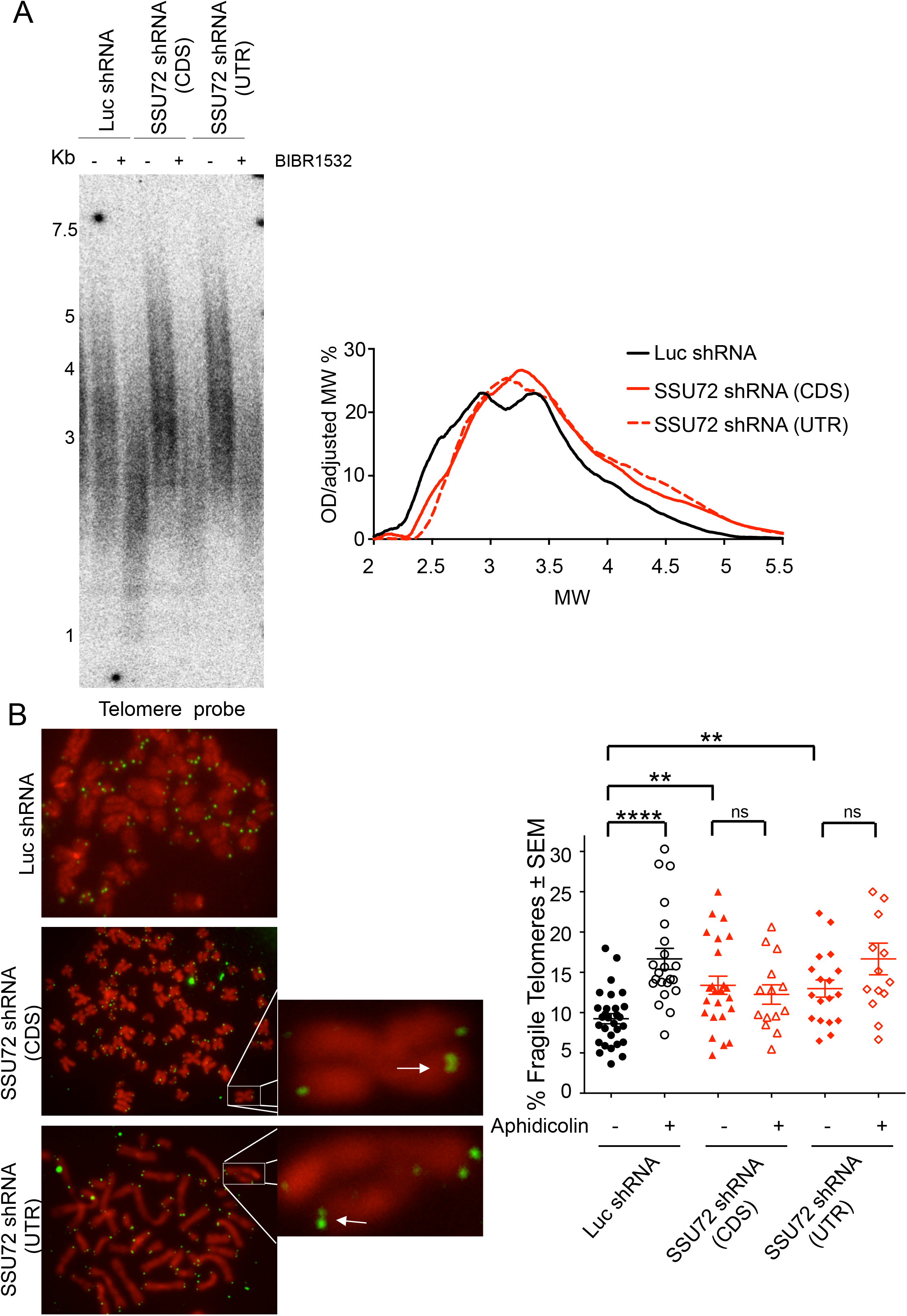
Down-regulation of human SSU72 results in telomere elongation and fragility. A) Telomere elongation of SSU72 down-regulated cells is telomerase dependent. HT1080 cells infected with lentiviral particles carrying two independent shRNAs against SSU72 (CDS and UTR regions) and control Luciferase (Luc) shRNA. Knockdown efficiencies were determined by RT-qPCR using specific primers against hSSU72. Quantification of Telomere restriction fragment analysis (TRFs) was carried out. B) hSSU72 down-regulation results in multi-telomeric signals (MTS) that are dependent on DNA replication. Visualisation of mitotic spreads of HT1080 hSSU72 shRNA cells treated with Aphidicolin and colcemid. FISH was carried out using a PNA-telomeric probe. Quantification of MTS: n=3; **p <0.01 *\*\*\*\*p* <0.0001 based on a two-tailed Student’s *t*-test to control sample. Error bars represent standard error of the mean (SEM).

We next asked if SSU72 downregulation results in DNA replication defects at telomeres, as is observed in fission yeast. As in previous studies, we used the appearance of multitelomeric signals (MTS) as a readout for faulty DNA replication at telomeres ^7^. Following shRNA treatment, we measured MTS in metaphase spreads of HT1080 cells and observed that, while 9.2% of telomeres showed MTS in control shLuciferase-treated cells, SSU72 downregulation using either CDS or UTR shRNA resulted in higher MTS levels (13.4% and 12.8%, p≤0.01) (**Figure 5B**). Previous studies have shown that treatment with Aphidicolin, a specific inhibitor of DNA polymerases, results in higher levels of MTS in mammalian cells ^7^. Indeed, Aphidicolin treatment of HT1080 cells resulted in elevated levels of MTS in control Luciferase shRNA-treated cells (p≤0.0001). In contrast, Aphidicolin did not result in increased MTS levels in cells where SSU72 had been downregulated using shRNAs against either CDS or UTR (p≤0.5 and p≤0.09, respectively) (**Figure 5B**). This result suggests that higher MTS levels observed in SSU72 downregulated cells are a consequence of collapsing replication forks, thus highlighting a role of SSU72 in controlling DNA replication in human cells. Consistent with our results, STN1 downregulation in human cells resulted in increased MTS levels ^41^. Moreover, STN1 dysfunction does not increase with Aphidicolin treatment, similar to findings in SSU72 downregulated cells ^41^. In parallel, we observed that the increase in MTS levels in SSU72 downregulated cells does not depend on telomerase activity. We observed that SSU72 downregulation is still able to induce higher MTS levels in U2OS telomerase-negative cells (Figure S6A). These data are consistent with STN1 dysfunction, as downregulation of this factor in U2OS cells induces equivalent rates of replication fork stalling at telomeres ^41^.

As expected, SSU72 downregulation in HT1080 cells also resulted in telomere induced foci (TIF), as measured by the localization of 53BP1 to telomeres (**Figure 6A**). Importantly, TIF formation was not cell line dependent, as we also observed TIFs in HeLa cells treated with an siRNA against SSU72 (**Figure S6B**).

**Figure 6.**
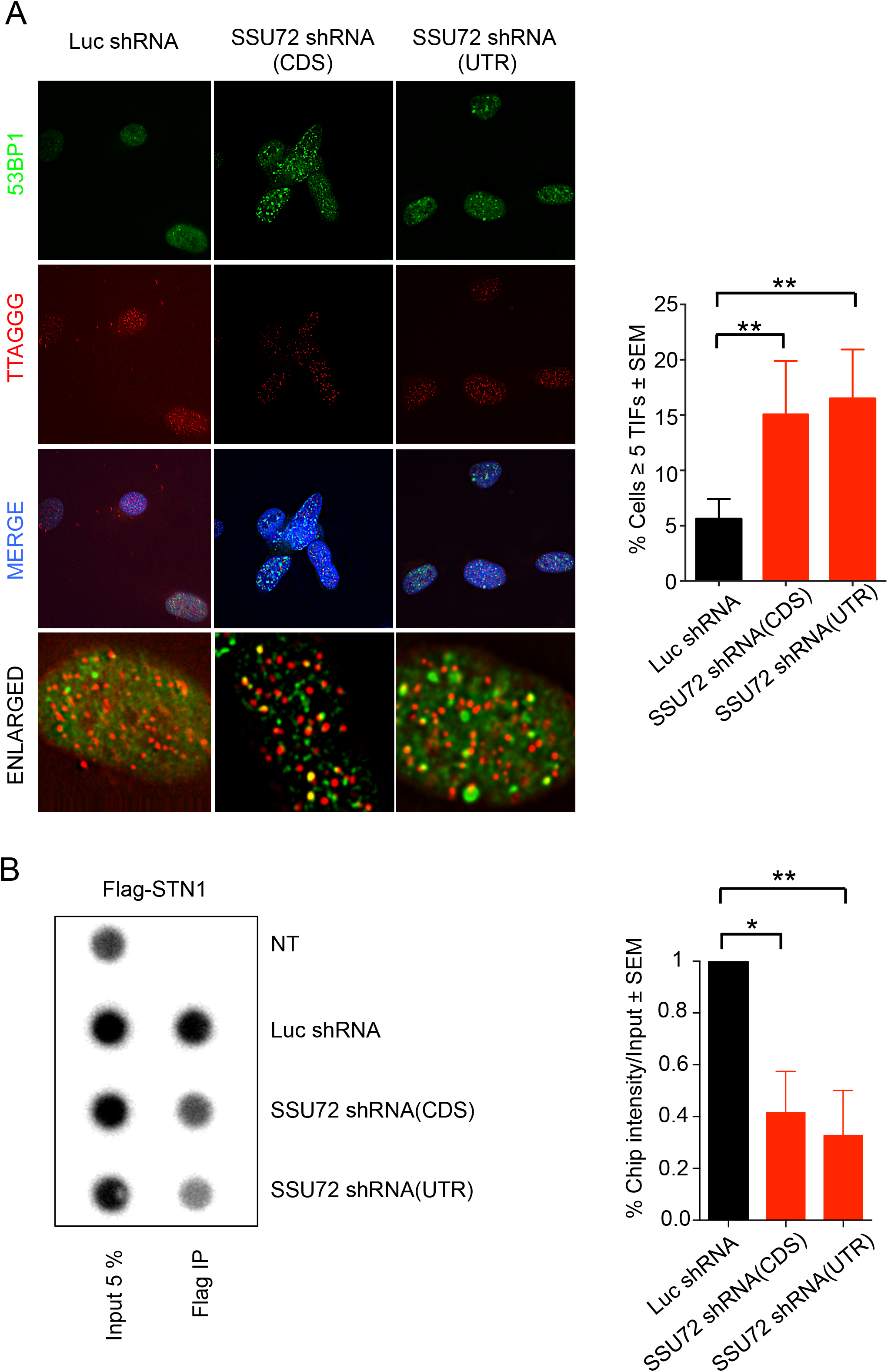
hSSU72 is required for hSTN1 recruitment to telomeres. A) hSSU72 downregulation results in telomere DNA damage foci (TIF). Cells with indicated shRNAs were fixed and IF-FISH was carried out using a 53BP1 antibody and PNA-telomere probes. Quantification of cells with more than 5 telomeric 53BP1 foci observed in A: n=3; **p <0.01 based on a two-tailed Student’s *t*-test to control sample. Error bars represent standard error of the mean (SEM). B) hSSU72 is required for efficient loading of hSTN1 at human telomeres. ChIP analysis was preformed using a FLAG antibody and Southern blotting was carried out using a human telomeric probe. Quantification of 3 independent ChIP experiments *p <0.05 **p <0.01 based on a two-tailed Student’s *t*-test to control sample. Error bars represent standard error of the mean (SEM).

Our data suggest that the downregulation of SSU72 in human cells mimics previous results obtained in STN1 downregulated cells. Thus, we tested whether cells lacking SSU72 were defective for STN1 recruitment to telomeres. We expressed FLAG-tagged STN1 in HT1080 cells and infected these cells with lentiviral particles expressing an shRNA against either SSU72 or Luciferase. We then carried out telomeric ChIP experiments using FLAG antibodies (**Figure 6B**). Upon downregulation of SSU72, we observed a 40% reduction in STN1 binding to telomeres compared to shLuciferase-treated cells. Together, these data indicate an evolutionarily conserved role for SSU72 phosphatase in controlling STN1 recruitment to telomeres and, therefore, in regulating lagging strand syntheses at telomeres.

## Discussion

Protein phosphorylation, a type of post-translational modification, plays key regulatory roles in almost all aspects of cell biology. Even though the function of protein kinases in telomere biology has been widely studied, the role of phosphatases remains relatively less explored. Contrary to this trend, budding yeast Pph22 phosphatase was recently shown to regulate the phosphorylation of Cdc13 in a cell cycle-dependent manner ^46^. The dephosphorylation of specific Cdc13 residues by Pph22 reverses the interaction of Cdc13 with Est1 and, consequently, disengages telomerase from telomeres ^46^. However, to date, there are no known phosphatases that regulate telomere length in fission yeast or higher eukaryotes. The data presented here depict an unprecedented role for a highly conserved phosphatase in telomere regulation. Ssu72 belongs to the group of class II cysteine-based phosphatases, which are similar to low molecular weight-phosphatases and some bacterial arsenate reductases ^47^. Although most well-known for its role as an RNA polymerase II CTD phosphatase in different species ^24^, human SSU72 has also been identified as a protein that can interact with the tumor suppressor Retinoblastoma ^48^ and can target STAT3 signaling and Th17 activation in autoimmune arthritis ^49^. Other functions have been reported in human cells, including SA2 dephosphorylation ^25^. Consistent with the multiple known roles of phosphatases, our work demonstrates that Ssu72 phosphatase also regulates telomere replication by controlling the recruitment of Stn1 to telomeres and promoting the Stn1-Pol1 interaction, thus activating lagging strand DNA synthesis.

At present, it is not well understood how lagging strand DNA replication inhibits telomerase activity. In fission yeast, Rad3 kinase is activated by the generation of ssDNA during DNA replication, leading to telomerase recruitment through the phosphorylation of Ccq1 at Thr93 ^27^. In this model, fill-in reactions by lagging strand polymerases reduce ssDNA at telomeres, thus contributing to a negative feedback loop. Consistent with the role of telomeric ssDNA in activating telomerase in fission yeast, *ssu72*Δ cells have longer overhangs, extensive phosphorylation of Ccq1 and higher levels of telomerase at telomeres. In addition, Rad3 and Ccq1-T93 phosphorylation are both required for the elongation of telomeres in *ssu72*Δ mutants. Therefore, it is still possible that Ssu72 phosphatase activity is required to regulate Ccq1 phosphorylation. Further experiments will be required to test this hypothesis.

Our work revealed that phosphorylation of Stn1 at Serine-74 is required for the regulation of telomere length. Mutation of Stn1 Serine-74 to Aspartic acid (D), an amino acid that mimics constitutive phosphorylation, results in telomeric elongation that is epistatic with the *ssu72*Δ mutation. This result indicates that Serine-74 phosphorylation is sufficient to explain the regulation of telomere length by Ssu72. The phosphorylation site identified in our mass spectrometry analysis resides within the OB fold domain of Stn1. OB fold phosphorylation is known to regulate protein-DNA binding and protein-protein interactions. For example, phosphorylation of human TPP1 (Tpz1 ortholog) in the OB fold domain regulates the telomerase-TPP1 interaction ^50^. In fission yeast, our data suggest that Stn1 phosphorylation at Serine 74 may prevent its binding to telomeric DNA in early S phase. Even though Serine 74 does not lie in a CDK consensus site, Cdc2^CDK1^ is likely to be the kinase responsible for Stn1 phosphorylation. Unlike other kinases, such as Hsk1^CDC7^, telomeric elongation in *ssu72*Δ mutants is reversed with increased inactivation of Cdc2 activity. Ssu72 counteracts Stn1 exclusion as it arrives at telomeres. Interestingly, Ssu72 is recruited to telomeres in the S/G2 phases concomitantly with the lagging strand machinery ^51^. Although further experiments are required, an attractive model involves Cdc2^CDK1^ phosphorylation of Stn1, thus creating a delay in lagging strand synthesis and allowing telomere elongation. Further, Ssu72 phosphatase reverses this process by promoting Stn1 binding to DNA polymerase alpha (**Figure 4E**) Notably, dephosphorylation of Stn1 has to be coordinated with Tpz1 SUMOylation, which is crucial for the recruitment of Stn1 to telomeres. Thus, both phospho- / dephosphorylation and SUMOylation-mediated interactions with Tpz1 control the recruitment and activity of telomeric Stn1-Ten1.

The SSU72 phosphatase appears to be conserved throughout evolution. The absence of a human SSU72 homolog in the HT1080 cell line results in similar phenotypes to those observed in fission yeast *ssu72*Δ mutants. First, SSU72 downregulation in HT1080 cells triggers DNA damage signaling at telomeres. Second, SSU72 depletion results in telomerase-dependent telomere elongation. Third, the recruitment of STN1 to telomeres is defective in SSU72-depleted HT1080 cells. Consistently, we observed increased telomere fragility (MTS) in SSU72-depleted cells, a phenotype also observed in STN1-deficient cells ^13^. We propose that SSU72 regulates STN1 recruitment to human telomeres in a manner similar to that in fission yeast. Currently, there is no evidence of STN1 phosphorylation in human cells. Nevertheless, Serine-74 is conserved in humans as an amino acid capable of being phosphorylated (T81). Further analysis will determine if human STN1 is phosphorylated at this residue and whether this modification regulates human telomere replication.

Recently, a model was proposed in which the replication fork regulates telomerase activity ^18^. This model describes how the regulation of origin firing and passage of the replication fork affect telomere homeostasis. In addition, we propose that telomere replication is controlled by two sets of phosphatases. On the one hand, the onset of telomere replication is regulated by Rif1-PP1A phosphatase through the inhibition of DDK activity at subtelomere origins of replication. On the other hand, we now show that telomere lagging strand synthesis is regulated by Ssu72 phosphatase, which promotes the Stn1-polymerase alpha interaction, thus terminating telomere replication and resulting in telomerase inhibition.

## Acknowledgments

We thank Dr. Nakamura, Dr. Nurse, Dr. Carr, Dr. Cooper and Dr. Bianchi for sharing their strains and Dr. Wang and Dr. Lingner for sharing their plasmids. We are grateful to Dr. Tomita, Dr. Jansen, Dr. Carlos, and Dr. Sridhar for reading the manuscript and to MGF laboratory for critical comments and discussions. This work was supported by the Portuguese Fundação Ciência e tecnologia (FCT) project number PTDC/BEX-BCM/5179/14. JME is supported by PTDC/BEX-BCM/5179/14 and PIEF-GA-2013-624759 and ESMC is supported by SFRH/BD/113754/2015. SM is supported by the Ligue Nationale Contre le Cancer (LNCC, Equipe Labellisée Vincent Géli). SC is supported by Projet Fondation ARC and by the Agence Nationale de la Recherche (ANR-16-CE12 TeloMito). IML acknowledges FCT for PhD fellowship funding (PD/BD/113982/2015) under the MolBioS PhD-Program (PD/00133/2012). IAA acknowledges IF/00764/2014 Research unit GREEN-IT “Bioresources for Sustainability” (UID/Multi/04551/2013). Mass spectrometry analyses were performed at UniMS (ITQB/iBET).

## Author Contributions

MGF and JME conceived the study and designed the experiments. JME and ESMC performed the majority of the experiments. MG and CR performed the fission yeast genetic screen. SC and SM performed the 2D gel experiments. IML and IA performed the mass spectrometry analysis. MGF and JME wrote the manuscript. MGF supervised the research.

## Material and methods

### Yeast strains and media

The strains used in this work are listed in *Supplementary Table 1*. Strains were constructed using commonly used techniques (Smith et al., 1999). Standard media and growth conditions were used throughout this work (Barinaga, 1997). For the strains containing pREP41 plasmids cultures were grown overnight in media PMG (Pombe Glutamate Medium) with the required amino acids. To generate the *ssu72-C13S*, ssu72+ gene was cloned into pGEM vector using genomic DNA. pGEM vector was mutagenized to create the pGEM-Ssu72-C13S. Endogenous ssu72+ gene was deleted using a *ura4+* fragment and FOA plates were used to select for Ssu72-C13S recombinants. Colonies were then screened for proper integration and sequenced to verify the presence of the point mutation. For *stn1-S74D* mutant strain, a genomic DNA fragment containing the *stn1+* gene was cloned into pJK210 plasmid and mutagenized to create the pJK210 *stn1-S74D*. The PacI-linearized pJK210 *stn1-S74D* plasmid was transformed into wild-type strain, and cells were plated on minimum medium lacking uracil. Ura4 positive cells were streaked on FOA-plate to select for direct-repeat recombination between *stn1+* and *stn1-S74D* allele. Presence of the *stn1-S74D* allele was subsequently verified by genomic sequencing.

### Southern blot analysis

Fission Yeast: Genomic DNA was obtained from exponentially growing yeast cells by phenol-chloroform extraction method. Human cells DNA was extracted as described in ^7^. Approximately 2 μg of digested *ApaI* or *EcoRI* DNA in Fission yeast or *Alul* and *Mbol* for human cells was run in either 1 % (fission yeast) or 0.6 % (human cells) agarose gels. The gel was transferred to a positively charged nylon membrane, and telomere analysis was performed as described (Rog et al., 2009) or (Reverter et al., 2010).

### Chromatin Immunoprecipitation (ChIP)

In Fission yeast, ChIP was performed as described (Moser et al., 2009). Briefly, exponentially growing cells were fixed with 1 % formaldehyde, 0.1M NaCl, 1mM EDTA, 50mM HEPES-KOH, pH 7.5 and Incubated 20 min at room temperature. Then, the solution was quenched with 0,25 M glycine (final concentration) for 5 minutes. After 2 washes with cold PBS, cells were lysed with 2 x lysis buffer (100 mM Hepes-KOH, pH 7.5 2 mM EDTA 2% Triton X-100 0.2% Na Deoxycholate) and disrupted by mechanical method. Chromatin was sheared, and equal amounts of DNA were used for immunoprecipitation protocol with either anti-Myc (9E10; Santa Cruz biotechnology) or anti-Flag (M2-F4802; Sigma) with magnetic Protein A beads. After washing the DNA-protein complexes with 1st (lysis buffer/0.1% SDS/275 mM NaCl), 2nd (lysis buffer/0.1% SDS/500 mM NaCl), 3rd (10 mM Tris-HCl, pH 8.0, 0.25 M LiCl, 1 mM EDTA, 0.5% NP-40, 0.5% Na Deoxycholate), Recovered DNA by 50 mM Tris-HCl, pH 7.5, 10mM EDTA, 1% SDS was decroslinked, purified and analysed in triplicate by SYBR-Green-based real-time PCR (Bio-Rad) using the primers described in ^52^.

For Human Chip, cells were fixed in 1% formaldehyde in PBS and incubated 15 min at room temperature. After quenching with Glycine, cells were lysed with 1% SDS, 50 mM Tris-HCl pH 8.0, 10 mM EDTA. After sonicating the chromatin, the diluted DNA protein complexes were incubated with Flag magnetic beads (Sigma M8823) overnight. After 3 consecutive washes, with 1x in IP buffer (20 mM Tris pH8, 0.15 M NaCl, 1% Triton X-100, 2 mM EDTA) with 0.1% SDS, 1x in IP buffer with 0.1% SDS, 0.5 M NaCl, and 10 mM Tris pH8.0, 1 mM EDTA with 1% Nonidet, 1% Na deoxycholate and 0.25 M LiCl. Complexes were eluted with 50 mM Tris pH 8.0, 10 mM EDTA, 1% SDS. Decrosslinked DNA was denatured and slot-blotted into a Hybond N+ membrane using a Bio-Rad blotter. Southern blot using a human telomere probe was carried out as described before ^53^.

### Immunoblotting

Whole-cell extracts prepared using exponentially growing yeast cells were processed for western blotting as previously described (Rog et al., 2009). Common western blot techniques were used to detect different proteins. For detection of Myc tagged proteins, we used anti-Myc monoclonal antibody (9E10; Santa Cruz) or rabbit anti-Myc (abcam). For detection of Flag-tagged proteins we used a Flag–M2 antibody (SIGMA - F1804).

### Immunoprecipitation

Exponentially growing yeast cells were lysed with IP Buffer x2 (50 mM HEPES [pH 7.5], 150 mM NaCl, 40 mM EDTA, triton 0.5% 0.1% NP40, 0.5 mM Na3VO4, 1 mM NaF, 2 mM PMSF, 2 mM benzamidine 10 % glycerol, Complete proteinase inhibitor + DNAse 10 u/ml). Equal amount of proteins was incubated with Flag–M2 (SIGMA - F1804) overnight followed by 3 washes for 10 minutes with IP buffer + 0.5 M NaCl. Common western blot techniques were then used to detect proteins.

### Mass spectrometry analysis

Stn1 protein was purified by tagging C-terminus with 13-myc tag. 5 litters of Logarithmic cycling cells were collected and lysed with 50 mM HEPES [pH 7.5], 150 mM NaCl, 40 mM EDTA, 0.2% Triton, 0.1% NP40, 0.5 mM Na3VO4, 1 mM NaF, 2 mM PMSF, 2 mM benzamidine, 10 % glycerol, DNAse I and Complete proteinase inhibitor. Cell lysates were incubated overnight with magnetic beads coated with Myc antibody (9E-10, Pierce). The immunoprecipitated was washed and run on a 4-12% Bis-Tris NuPAGE gel (Invitrogen). A slice of gel ranging between 75 and 63 kDa was excised and tryptic peptides were prepared by *in-gel* digestion ^54^. Peptides were analyzed by nanoLC-MS using an Ekspert 425 nanoLC with cHiPLC (Eksigent, AB Sciex, Framingham, MA USA) coupled to a TripleTOF^™^ 6600 mass spectrometer (AB Sciex, Framingham, MA, USA).

Spectra were searched against Swiss-Prot database (downloaded in 10/2017, 5201 entries) containing all the reviewed protein sequences available for S. *pombe*, and three human keratin sequences (P04264, P35908, P13645). The Paragon algorithm embedded in ProteinPilot 5.0 software (AB Sciex, Framingham, MA USA) was used to perform the database search using phosphorylation emphasis and gel based ID as special factors, and biological modifications as ID focus. An independent false discovery rate (FDR) analysis was carried out using the target-decoy approach provided with Protein Pilot software and positive identifications were achieved using a global FDR threshold below 1%.

## Two-dimensional (2D) gel electrophoresis

2D gel electrophoresis experiments were carried out as described in ^55^. 10 μg of DNA (for telomeres analysis) was digested with 60U of *NsiI*. For analysis of the RFB region, 5 μg of DNA was digested with 60 U of *BamHI*. DNA was run on 0.4% agarose gel for the first dimension and a 1% agarose gel for the second dimension. Gels were transferred to positively charged membranes and probed with the STE1 probe or the 1.35-kb *EcoRI-EcoRI* rDNA fragment.

## Human lentiviral infections

HT1080 cell line were infected with either Luciferase shRNA (target sequence CGCTGAGTACTTCGAAATGTC), CDS shRNA (target sequence CAAAGACCTGTTTGATCTGAT) or UTR shRNA (target sequence ACGGTAGCATTACCCAAATAA) lentiviral particles produced in 293T cells by mixing PLKO vectors with psPax2 and pVSVG vectors. Cells were infected twice and selected with puromycin at 3 micrograms/ml.

### Metaphase spreads

Metaphases were collected by adding Colcemid to the media to a 0.1μg/ml final concentration for 4 hours to overnight. Metaphases were collected using the shake off method. Cells were then incubated in hypotonic buffer (0.03M Na citrate) for 30 minutes and fixed with Methanol:acetic acid (3:1) solution. Slides were pre-washed with 45% acetic acid spread metaphases.

### Fluorescence In Situ Hybridization (FISH)

For FISH using telomere probes, slides were washed in PBS 3×5min each with rotation and incubated with 10 mM Tris, pH7.5 and a deionized formamide 70% telomere probe (0.5 ng/ml), blocking reagent (0.25%, 25 mM MgCl2, 9mM citric acid, 82mM Na2HPO4, pH7.0) at 80°C for 3 minutes. Slides were then enclosed in a humidifier chamber for 3 hours. After serial washes of 70% formamide, 10 mM Tris, 0. 1% BSA (2 times for 15 minutes) and PBS-Tween 0.05% (3 times for 5 minutes), slides were dried and mounted with mounting media with 1 μg/ml DAPI. For slides with metaphase spreads a first step of 37 % pepsin digestion (0.1 g/100 ml + 88 μl HCl) before starting the FISH technique

### Microscopy

In fission yeast, cells were grown at 32°C in PMG with all supplements added. For each individual experiment, at least 100 cells were analysed. For GFP and mRFP visualization, live cells were imaged using a Delta Vision Core System (Applied Precision) using a 100× 1.4 numerical aperture UplanSApo objective and a cascade2 EMCCD camera (Photometrics). Deconvolution was performed using the enhanced ratio method in softWoRx software. Co-localization experiments were performed using maximum intensity projections of deconvolved images.

For Immunofluorescence-FISH analysis in human cells, cells were fixed in 2-4% formaldehyde in PBS for 10 min at RT. After SDS (0.03%) permeabilization and 15 min blocking step (1% BSA, 0.5% Triton X-100, 0.5% Tween 20), 53PB1 antibody (1:1000; Santa Cruz biotechnology H-300) was incubated overnight. Antibody was washed 3 times with PBS-Tween 0.05% and a secondary antibody conjugated with alexa 488 (1:400) was used ^56^. Before starting the FISH technique, fixation of the cells was performed using 2-4 % formaldehyde for 10 min at room temperature.

---

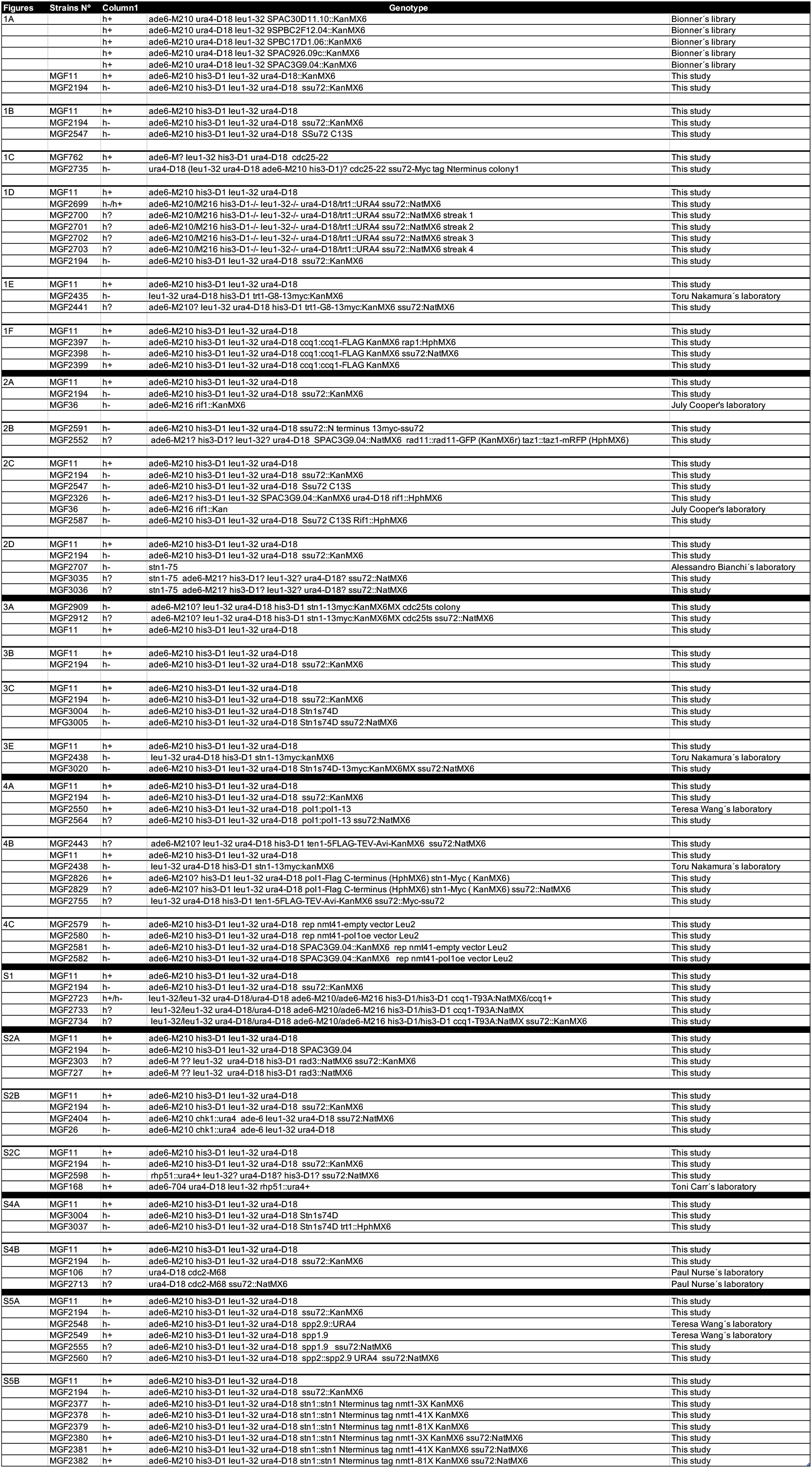
Supplementary table 1. List of strains used in this manucript

**Figure S1.**
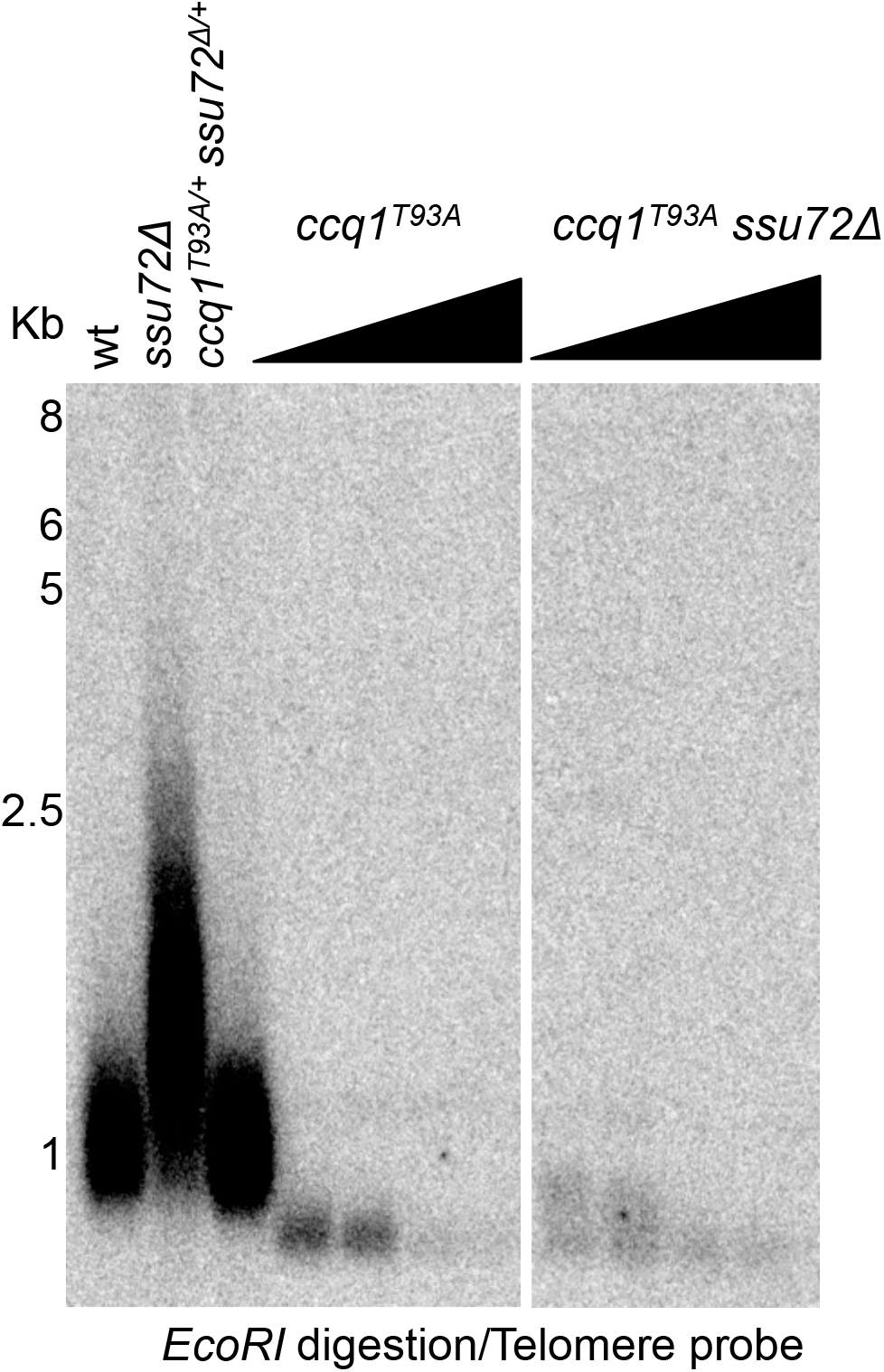
Phosphorylation of Ccq1 in Threonine93 is required for telomere elongation in *ssu72* mutant. Diploid strains with the appropriate phenotypes were sporulated and streaked for different passages. Telomere length was measured in *EcoRI* digested DNA by a telomeric probe.

**Figure S2.**
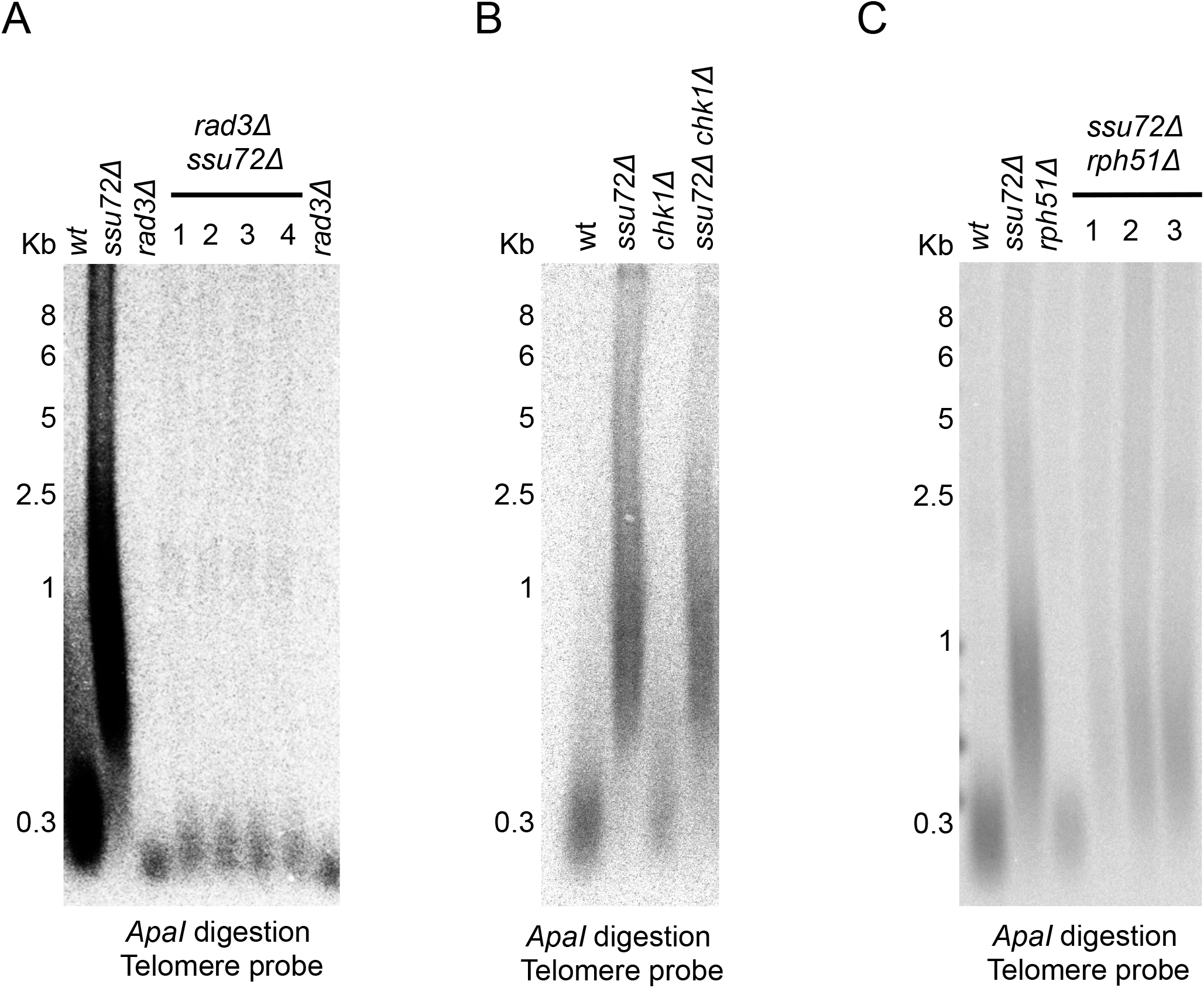
Telomere length in *ssu72Δ* is *rad3*Δ (A) dependent, but checkpoint (B) and homologous recombination (C) independent. *rad3Δ(A) chk1*Δ (B), *rph51*Δ (C) single mutants or different colonies of double mutants *ssu72Δ-rad3Δ (A), ssu72Δ-chk1Δ (B), ssu72A-φh51Δ (C)*were constructed and telomere length was measured carrying out Southern blots in *Apal* digested genomic DNA using a telomeric probe.

**Figure S3.**
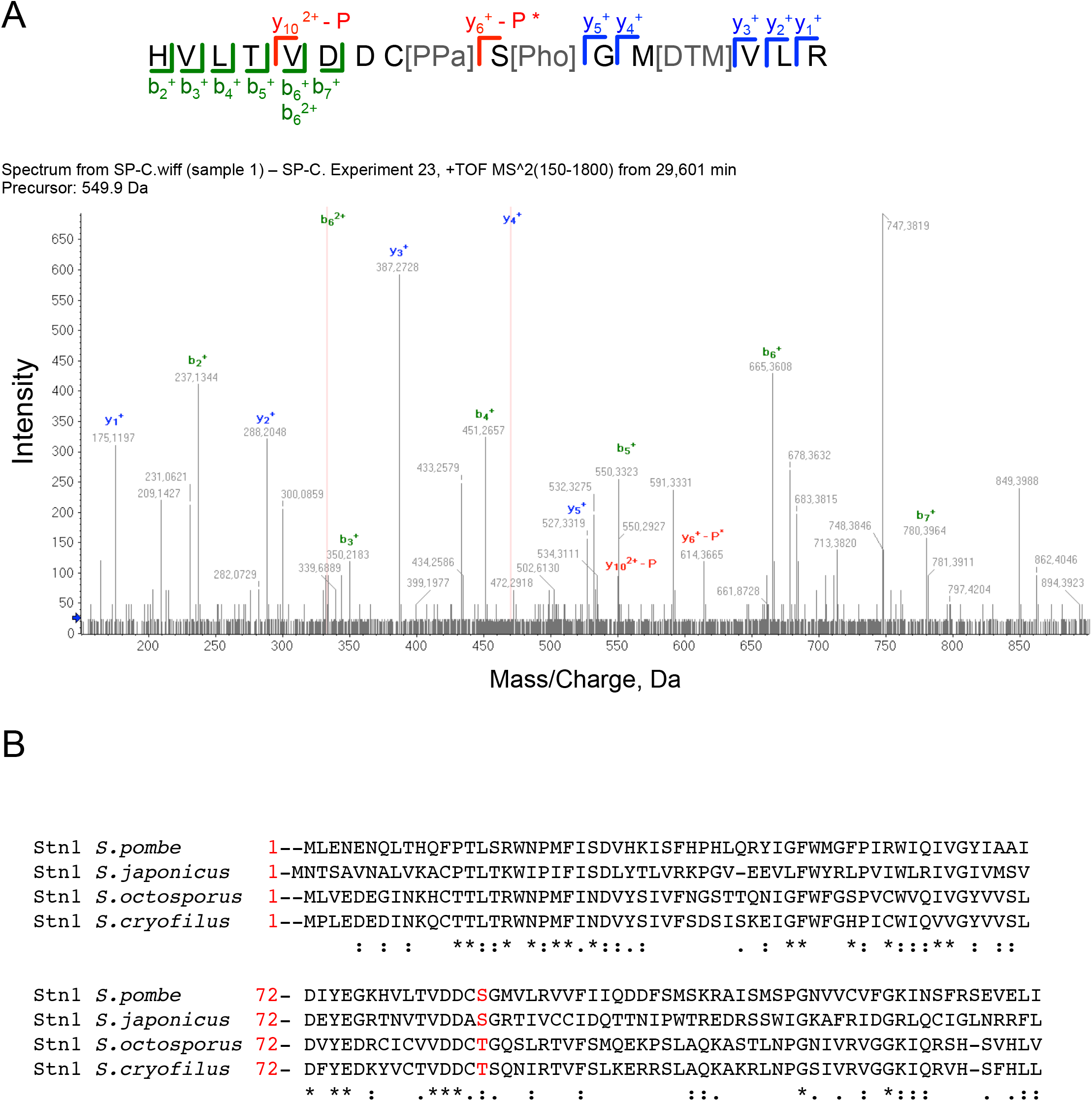
A) Identification of S74 as Stn1 phosphorylation site in fission yeast. Mass spectrometry spectra identifying the phosphorylated peptide on the Stn1-S74 residue **B) S74 residue is conserved in Schizosaccharomyces family.** Sequence alignment of Stn1 protein in *Schizosaccharomyces* family (*S. pombe, S. cryophilus, S. octosporus, and S. japonicus*) using Clustal Omega. Serine identified in fission yeast as phosphorylated is squared in red. The site is either conserved or substituted by other aminoacid that is capable to be phosphorylated in Schizosaccharomices family.

**Figure S4.**
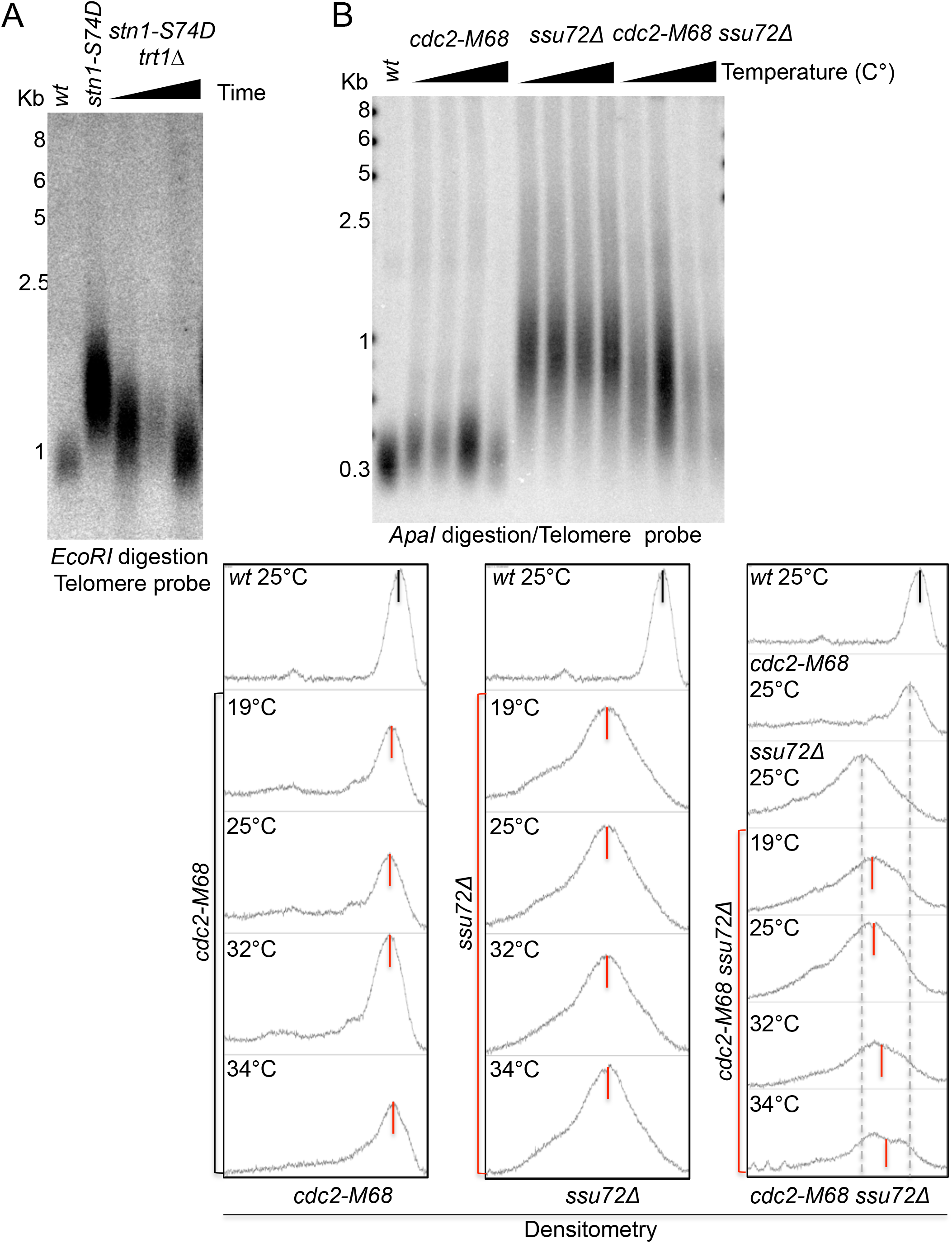
A) Telomere length of *stn1-S74D* is dependent on telomerase. *trt1^+^* was deleted in the *stn1-S74D* background and double mutants were streaked for multiple passages (triangle indicates increased number of generations). **B) Cdk1 activities are required to elongate telomeres in a *ssu72*Δ background.** Cells were grown at 25°C and then shifted to different temperatures by 16 hours to partially inactivate cdc2. DNA was isolated and telomere length measured with *Apal* digested DNA. Strains were constructed by regular techniques. B) Telomere length was measured with ImageJ. Black and Red lines represent average telomere length in wt, cdc2-M68, *ssu72*Δ or double mutant cdc2-M68 *ssu72*Δ strains.

**Figure S5.**
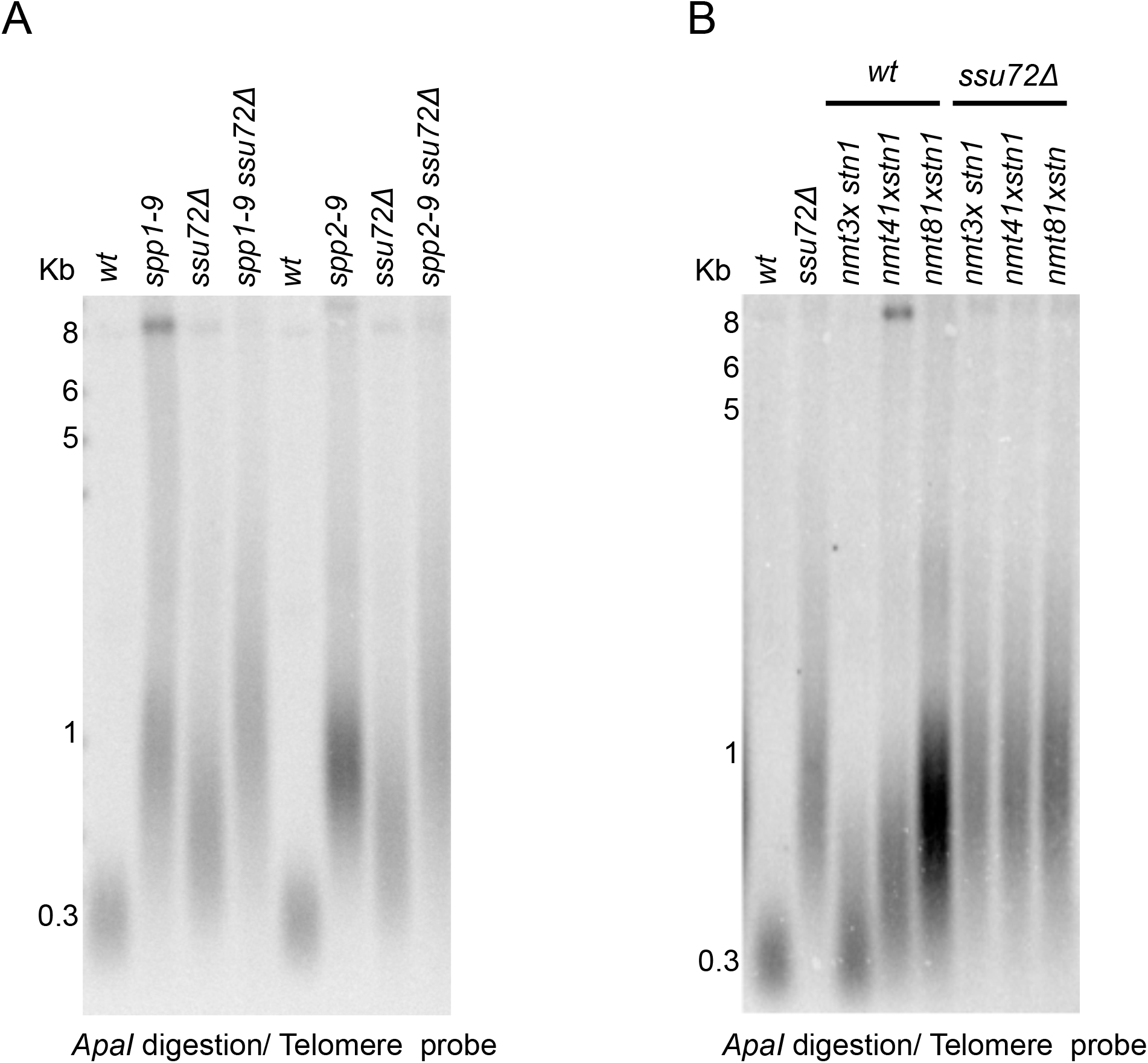
**A) *ssu72*Δ telomere length is epistatic with polymerase alpha complex subunits** Genomic DNA of single mutants of *pol1-13, spp1-9, spp2-9, ssu72Δ* and *wt or* double mutants *spp1-9 ssu72Δ* and *spp2-9 ssu72Δ* was isolated and telomere length was measured carrying out a Southern blot in *Apal* digested genomic DNA using a telomeric probe. Temperature sensitive strains were grown at semi-permissive temperature by several generations and DNA was collected to carry out Southern Blot analysis. **B) Stn1 overexpression doesn’t rescue telomere defect in *ssu72*Δ**. We expressed *stn1* under 3x (stronger), 41 x and 81 x (weaker) nmt1 promoter in wŕ or *ssu72*Δ background and telomere length was measured in *Apal* digested genomic DNA.

**Figure S6.**
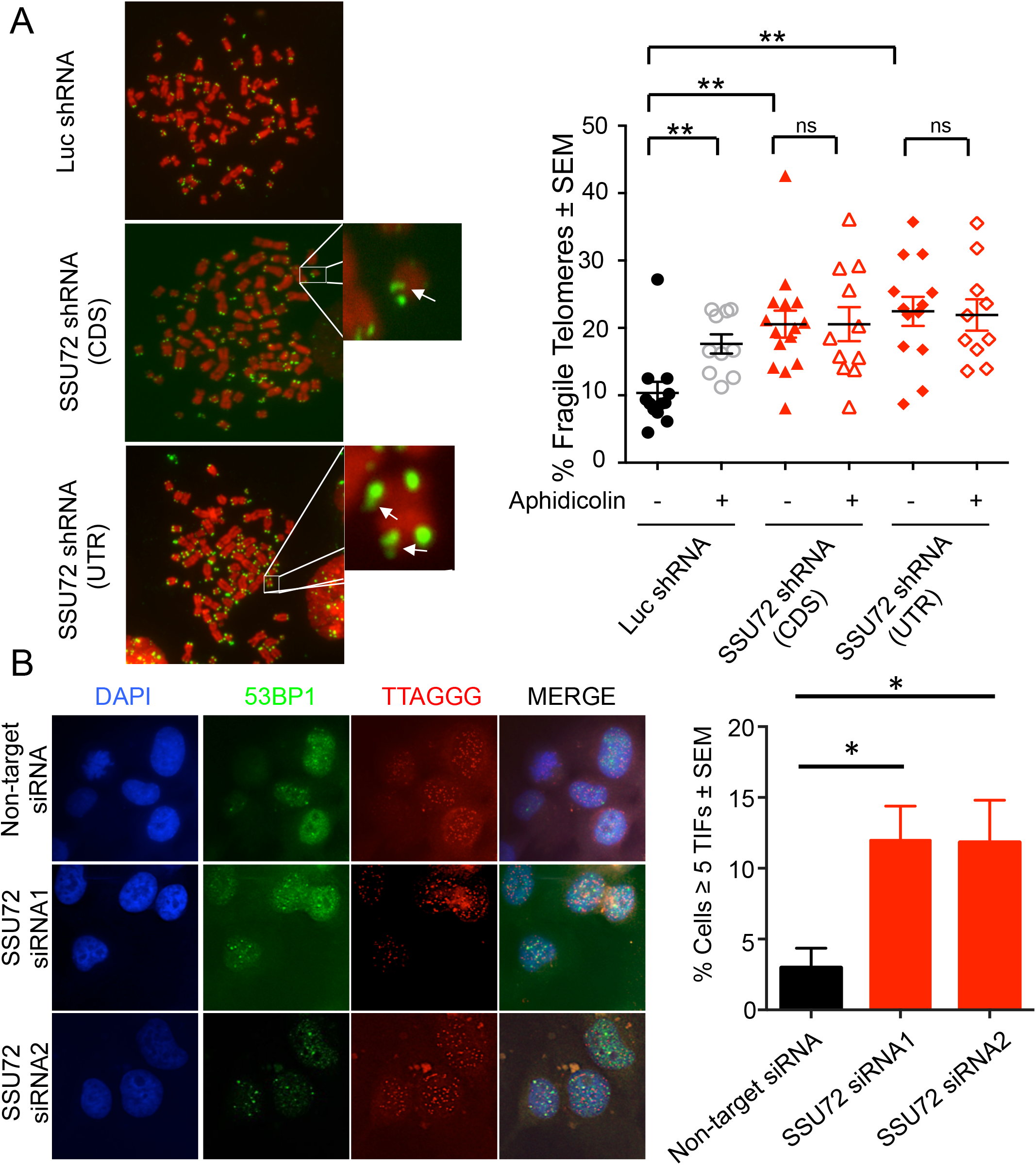
A) SSU72 downregulation induces telomere fragility in U20S cell line. Cells were infected with lentiviral particles carrying shRNAs against either SSU72 or Luciferase shRNAs. If appropriate, cells were treated with Aphidicolin at 200 ng/ml for 12 hours before colcemid treatment. Metaphases were collected and FISH was carried out using a PNA- telomeric probe. Quantification of MTS in SSU72 downregulated cell was carried out n = 2; **p <0.01 based on a two-tailed Student’s *t*-test to control sample. Error bars represent mean ±standard error of the mean. **B) SSU72 downregulation induces 53BP1 foci at telomeres in Hela human cell line.** Cells were transfected with two independent s¡RNAs against human SSU72 using a non-targeting s¡RNA as a control. After 3 days cells were fixed and IF-FISH was carried out using a 53BP1 antibody and PNA- telomeric probe. Quantification of Telomeric induce foci (TIF) in SSU72 downregulated cell was carried out n = 3; *p <0.05 based on a two-tailed Student’s *t*-test to control sample. Error bars represent mean ±standard error of the mean.

